# Engineering asymmetric nanoscale lipid vesicles for drug delivery

**DOI:** 10.1101/2024.08.30.610290

**Authors:** Chenjing Yang, Julian Menge, Nene Zhvania, Dong Chen, David A. Weitz, Kevin Jahnke

## Abstract

The delivery of therapeutics to cells enables both the treatment and the prevention of diseases. To protect therapeutics from degradation and enable cell-specific targeting, they are often encapsulated into drug delivery vehicles such as lipid nanoparticles, viral vectors or lipid vesicles. These delivery vehicles have been extremely successful in delivering small molecules, nucleic acids or proteins. However, there is no universal drug delivery vehicle that can deliver therapeutics irrespective of the choice of cargo. Here, we present a method to engineer lipid vesicles with asymmetric leaflets and show that they can deliver mRNA and proteins to cells. We also find that the leaflet asymmetry can increase the lipid vesicle uptake by cells. When we load asymmetric vesicles with mRNA, we observe a 5-fold increase in the transfection efficiency indicative of an improved uptake and release by asymmetric vesicles. Moreover, our findings extend beyond mRNA cargos by showcasing the effectiveness of asymmetric vesicles in delivering a wide range of proteins to cells, including the promising CRISPR/Cas9 gene editing system. Our method and findings expand the parameter space for engineering drug delivery vehicles and demonstrate the pivotal role of leaflet asymmetry in determining the performance of drug delivery vehicles. Consequently, our work leads to many applications, including the formation of more efficient universal drug carriers and the delivery of gene-editing proteins to cells.

## Introduction

To prevent the degradation of therapeutics and clearance by the immune system, drugs are often packaged into delivery vehicles.^1–3^ Following the uptake by cells, these drug delivery vehicles release the drug into the cytoplasm.^4,5^ The drugs can vary widely in size and their biochemical properties, ranging from relatively simple small molecules to nucleic acids and even proteins such as the gene-editing complex CRISPR/Cas9.^1,6,7^ As drugs and therapeutics have become more complex, so have their delivery vehicles, due to the increasing challenges posed by the large sizes and intricate charge properties of modern drugs.^8^ Lipid nanoparticles, which have been prominently used for the COVID19 vaccines, are extremely successful because they can encapsulate negatively charged nucleic acids such as mRNA.^9,10^ The electrostatic interactions of positively charged lipids and negatively charged cargo leads to the formation of nanoparticles with high encapsulation efficiencies.^11^ However, lipid nanoparticles still face limitations in delivering neutral or positively charged therapeutics.^12^ Additionally, a change in the lipid composition will invariably affect the encapsulation efficiency and lipid nanoparticle size.^13^ This can be particularly important for the encapsulation and delivery of more complex cargos such as proteins.^14^ Proteins are larger than nucleic acids and have an intricate charge profile, which make them difficult to encapsulate into lipid nanoparticles. By contrast, viral vectors are an alternate type of delivery vehicle and do not rely on electrostatic interactions to form. Instead, they use natural virus capsids as compartment to host protein cargo. They are very promising for the delivery of proteins and have been used to efficiently transfect cells.^15^ However, they are difficult to produce in large-scales and are often immunogenic, leading to an elevated immune response and clearance by the liver.^16^ Moreover, their size is set by the virus capsid and cannot be easily tuned causing many viral vectors to either be too small for the efficient loading with proteins or too large to transfect cells.^17^ Lipid vesicles are a third compartment type formed by amphiphilic lipid molecule bilayers.^18–20^ They can be assembled at high concentrations and in large quantities, while having complete control over their size which can range from 30 nm to 100 µm.^20^ Even though their charge can be tuned through the lipid composition, the encapsulation of therapeutics is challenging because the cargo is typically loaded after the vesicle has formed.^21^ Therefore, lipid vesicles are currently only used for small molecules that can permeate the membrane upon chemical manipulation.^22^ This has been implemented in 14 US Food and Drug Administration (FDA)-approved treatments for small molecules but the delivery of more complex cargo such as mRNA or proteins is difficult.^23^ Thus, while all delivery vehicles have important applications, none can simultaneously meet all the requirements for universal cargo delivery, such as encapsulating arbitrary macromolecular cargo and maintaining size control.^24^ If we could develop a strategy to encapsulate larger cargo into lipid vesicles and overcome their low uptake by and release in cells, we would enable the treatment of many diseases that are currently out of the scope of conventional delivery vehicles.

Here, we engineer a lipid vesicle capable of encapsulating and transfecting cells with therapeutics, including proteins and nucleic acids. To achieve this, we assemble nanoscale lipid vesicles with asymmetric leaflets using an inverted emulsion technique. We extrude water-in-oil emulsions to make them sufficiently small and centrifuge them through a second lipid monolayer to form small asymmetric lipid vesicles. We verify the leaflet asymmetry of up to 93% using fluorescence quenching experiments. Remarkably, we find that leaflet asymmetry can improve lipid vesicle uptake by cells, the mRNA transfection efficiency, reduce cell cytotoxicity and deliver a wide range of proteins to cells, including the gene-editing protein Cas9. The uniqueness of asymmetric lipid vesicles broadens the parameter space for the engineering of drug delivery vehicles and leads to many potential applications, including the formation of more efficient drug carriers with reduced toxicity while also enabling new treatment options that require the delivery of proteins.

## Results & Discussion

We engineer nanoscale lipid vesicles with asymmetric leaflets using the inverted emulsion technique coupled to an extrusion step. We form an inverted emulsion consisting of water droplets dispersed in mineral oil, which contains lipids as surfactants.^25–28^ The lipids form a monolayer that stabilizes the water-in-oil droplet and serves as the inner leaflet of the lipid vesicle.^29,30^ Subsequently, we extrude the water in mineral oil emulsions through a polycarbonate membrane with holes of a fixed diameter to generate monodisperse nanoscale emulsions. These emulsions are then centrifuged through a second lipid monolayer at a second water/oil interface to produce asymmetric lipid vesicles, schematized in Figure 1**a**. This method yields asymmetric vesicles because both monolayers form independently as they are spatially separated until the centrifugation step. We can adjust the diameter of asymmetric lipid vesicles by using polycarbonate membranes with different pore sizes. With a 200 nm membrane, the hydrodynamic vesicle diameter is about 250 nm; by using a 100 nm membrane we reduce the vesicle size to 150 nm, whereas a 30 nm membrane yields vesicles with a hydrodynamic diameter of 65 nm, as shown in Figure 1**b**. In addition to the scattering experiments, we also validate the vesicle size using transmission electron microscopy (TEM) in Figure S1. A key advantage of this method is that it yields asymmetric vesicles. We measure the degree of asymmetry with a dithionite quenching assay, where the outer leaflet is labeled with an NBD-PC fluorescent probe. NBD is quenched upon exposure with membrane impermeable dithionite, allowing us to quantify the amount of lipids that has flip-flopped to the inner leaflet. We observe that the fluorescent intensity decreases to approximately 10% after the addition of the dithionite and further reduces to 1% after the addition of Triton X, which destroys the lipid vesicles. This confirms that we form lipid vesicles with 90% asymmetry in their lipid composition. We prove the asymmetric properties of many different lipid compositions, as shown in Figure S2 and verify that the lipid asymmetry remains even after 24 h, as illustrated in Figure 1**c**. To confirm that we can tailor the composition and charge of lipids in both the inner and outer leaflet, we measure the zeta potential of lipid vesicles with different ratios of positively and negatively charged lipids. By keeping the inner leaflet containing uncharged POPC lipids fixed and changing the amount of positively charged EPC lipids in the outer leaflet, we find that the zeta potential is increasing proportionally to the amount of positively charged lipids up to 50 mV. This value is consistent with symmetric lipid vesicles made purely from EPC with extrusion as shown in Figure S3. When we replace EPC lipids with negatively charged POPS, we observe a reduction of the zeta potential to -50 mV as shown in Figure 1**d**. By contrast, when we keep the outer leaflet uncharged and increase the ratio of the charged lipids in the inner leaflet, the zeta potential remains constant, as shown in Figure 1**e**. This shows that the zeta potential is only influenced by the lipids in the outer leaflet.

**Figure 1:**
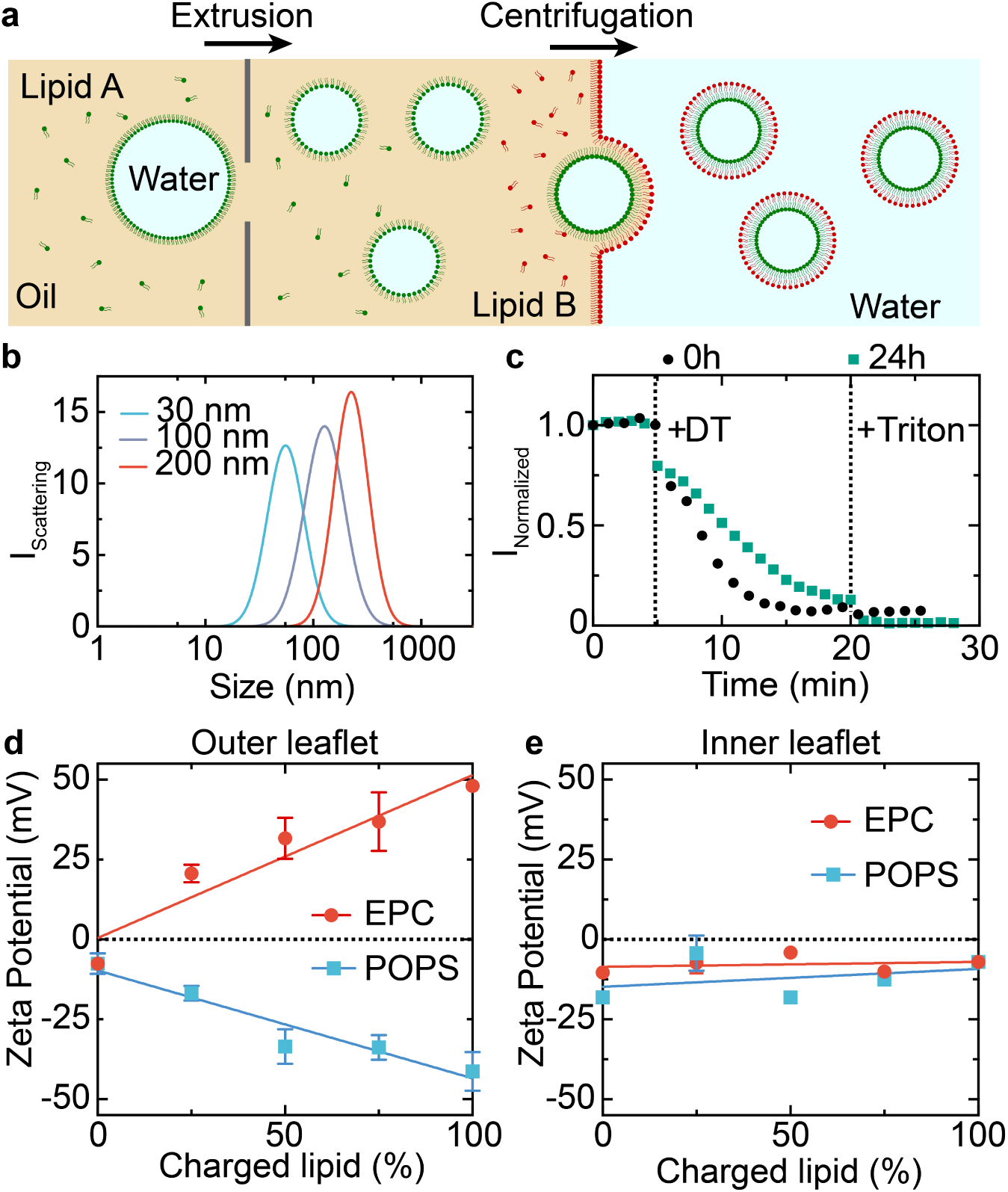
Engineering small asymmetric vesicles via an inverted emulsion coupled to an extrusion step. **a** Schematic representation of the process used to form small asymmetric vesicles. **b** Scattering intensity of small asymmetric vesicles for different polycarbonate membranes used for the extrusion. **c** Normalized fluorescence intensity of asymmetric lipid vesicles during the dithionite quenching assay with NBD-labeled lipids in the outer leaflet. **d** Zeta potential of asymmetric vesicles as a function of the EPC and POPS content in the outer leaflet. **e** Zeta potential of asymmetric vesicles as a function of the EPC and POPS content in the inner leaflet. Mean *±* SD, n=3.

The properties of lipid vesicles critically depend on the lipid composition and charge. To determine if we can assemble a wide variety of asymmetric lipid vesicles, we test the vesicle formation with four different charged lipid compositions: uncharged POPC, negatively charged POPS and positively charged DODMA and EPC as shown in Figure S4. Strikingly, we managed to form vesicles for all permutations of lipids in each leaflet. This even includes the formation of vesicles with oppositely charged lipids in each leaflet. However, we observe that each lipid composition has a different yield of vesicles. We measure the yield of lipid vesicles by adding a small amount of a fluorescently labelled lipid to each leaflet. Thus, the fluorescence intensity is directly proportional to the yield of lipid vesicles. We observe that the yield is always highest when POPS is in the outer leaflet irrespective of the inner leaflet as shown in Figure 2**a**. When the outer leaflet is composed of POPC, the yield is reduced by more than 3-fold and even more when we use EPC or DODMA.

**Figure 2:**
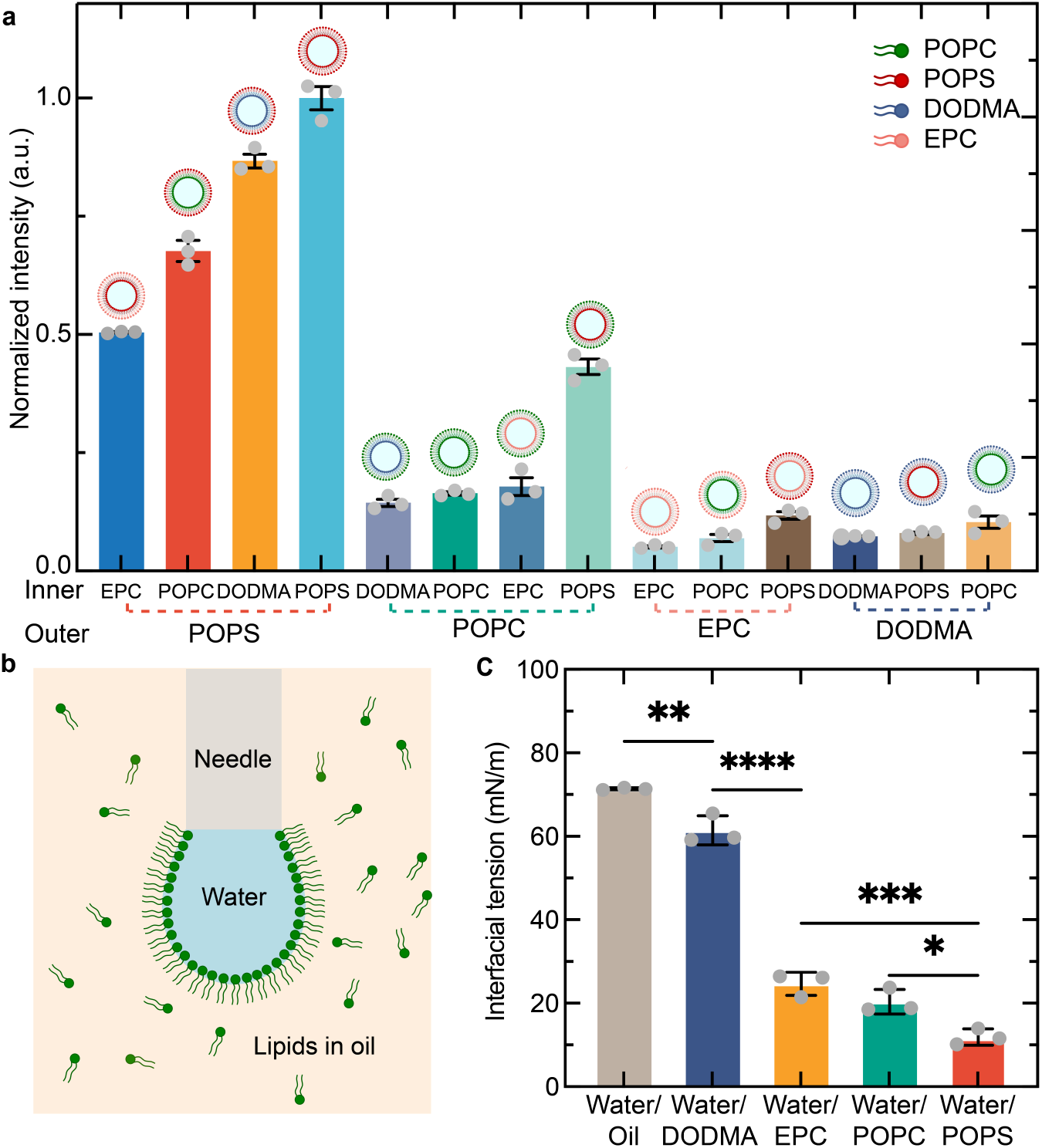
The yield of vesicles depends on the choice of lipids. **a** Fluorescence intensity correlating with the vesicle yield for lipid vesicles formed with DODMA, EPC, POPC and POPS (inner-outer leaflet). **b** Schematic representation of the pendant drop method used to measure the interfacial tension between water and different mineral oil solutions. **c** Interfacial tension between different water/mineral oil formulations in presence of DODMA, EPC, POPC, POPS or in the absence of lipids. Mean *±* SD, n=3.

During the formation of asymmetric lipid vesicles, it is crucial for the lipid-stabilizing emulsions to translocate from the oil phase to the water phase. For this reason, we investigate the effect of lipids at the water/oil interface by measuring the interfacial tension with the pendant drop method, as schematized in Figure 2**b**. The interfacial tension results show that DODMA has the highest interfacial tension with 61.5*±*3.5 mN/m. By contrast, EPC (25 mN/m)significantly reduces the interfacial tension followed by POPC (20 mN/m) and POPS (12 mN/m), as shown in Figure 2**c**. This reveals an interesting correlation between the yield of vesicles and the interfacial tension at the second water/oil interface. The lower the interfacial tension, the higher the yield of vesicles. We also find that lipids that reduce the interfacial tension lead to a larger amount of partitioning of hydrophilic molecules from the water to the aqueous phase, as shown in Figure S5. The relationship between interfacial tension and lipid vesicles yield suggests that the addition of co-surfactants could increase the yield of lipid vesicles.

To investigate the potential of asymmetric lipid vesicles as drug delivery vehicles, we label them with a fluorescent lipid and add them to human embryonic kidney (HEK) cells. We calibrate the concentration of the vesicles added to the cells by comparing the yield with extruded lipid vesicles, as shown in Figure S6 and Note S1. We always add the same lipid concentration (approximately 25 µM) of lipid vesicles to HEK cells and observe the lipid vesicle uptake after 24 hours of incubation. The confocal images show that the cells barely take up POPC-POPC (inner-outer leaflet) vesicles. By contrast, the uptake is significantly higher for POPS-POPS, as shown in Figure 3**a**. When we quantify the lipid vesicle uptake per cell from confocal images, we also observe a higher uptake of POPS-POPS vesicles compared to POPC-POPC vesicles. We confirm this with symmetric extruded lipid vesicles as shown in Figure S7. Thus, the incorporation of PS in lipid vesicles enhances the cell uptake. To quantify a larger number of cells, we also use flow cytometry to analyze the lipid vesicle uptake per cell. Unexpectedly, we observe a 2-times higher lipid vesicle uptake for asymmetric POPC-POPS lipid vesicles compared to symmetric POPS-POPS vesicles as shown in Figures 3**a**, 3**b** and Figure S8. This reveals that not only the outer leaflet, which directly interacts with the cell membrane, dictates the lipid vesicle uptake but also the lipids in the inner leaflet. The uptake of asymmetric POPS-POPC vesicles with the inverted lipid composition is 4 times lower than the uptake of POPC-POPS. To test by which mechanism the lipid vesicles are taken up by cells, we investigate the lipid vesicle uptake for cells while inhibiting a specific endocytosis pathway. As shown in Figure 3**c**, the lipid vesicle uptake decreases 3-fold when we add cytochalasin D and dynasore, two drugs that inhibit the clathrin-mediated endocytosis and macropinocytosis pathways. This is in good agreement with the endocytosis mechanisms for lipid nanoparticles and conventional lipid vesicles.^31^ We further verify these flow cytometry results with confocal microscopy as shown in Figure S9. To explain the difference in the uptake by cells, we hypothesize that the inner leaflet changes the biophysical properties of the asymmetric lipid vesicles such that they are softer than their symmetric counterparts. Soft lipid vesicles deform easily during endocytosis, increase the contact area between lipid vesicles and the cell membrane and hence decrease the energy barrier required for cells to take up lipid vesicles. To test this hypothesis, we prepare vesicles with different stiffness by changing the length of the fatty acids. When we add lipid vesicles containing 18:0 PS lipid vesicles, the cell uptake is the much smaller than for 16:0 PS (POPS) lipid vesicles. By contrast, the uptake for 15:0 PS lipid vesicles is much higher than for POPS, as shown in Figure S10. This shows that the cell uptake increases for vesicles made from lipids with shorter fatty acids, which likely correlates with softer vesicles. These results indicate that vesicle stiffness, in addition to the choice of lipids on the outer leaflet, is an important factor that drives the lipid vesicle uptake by cells.

**Figure 3:**
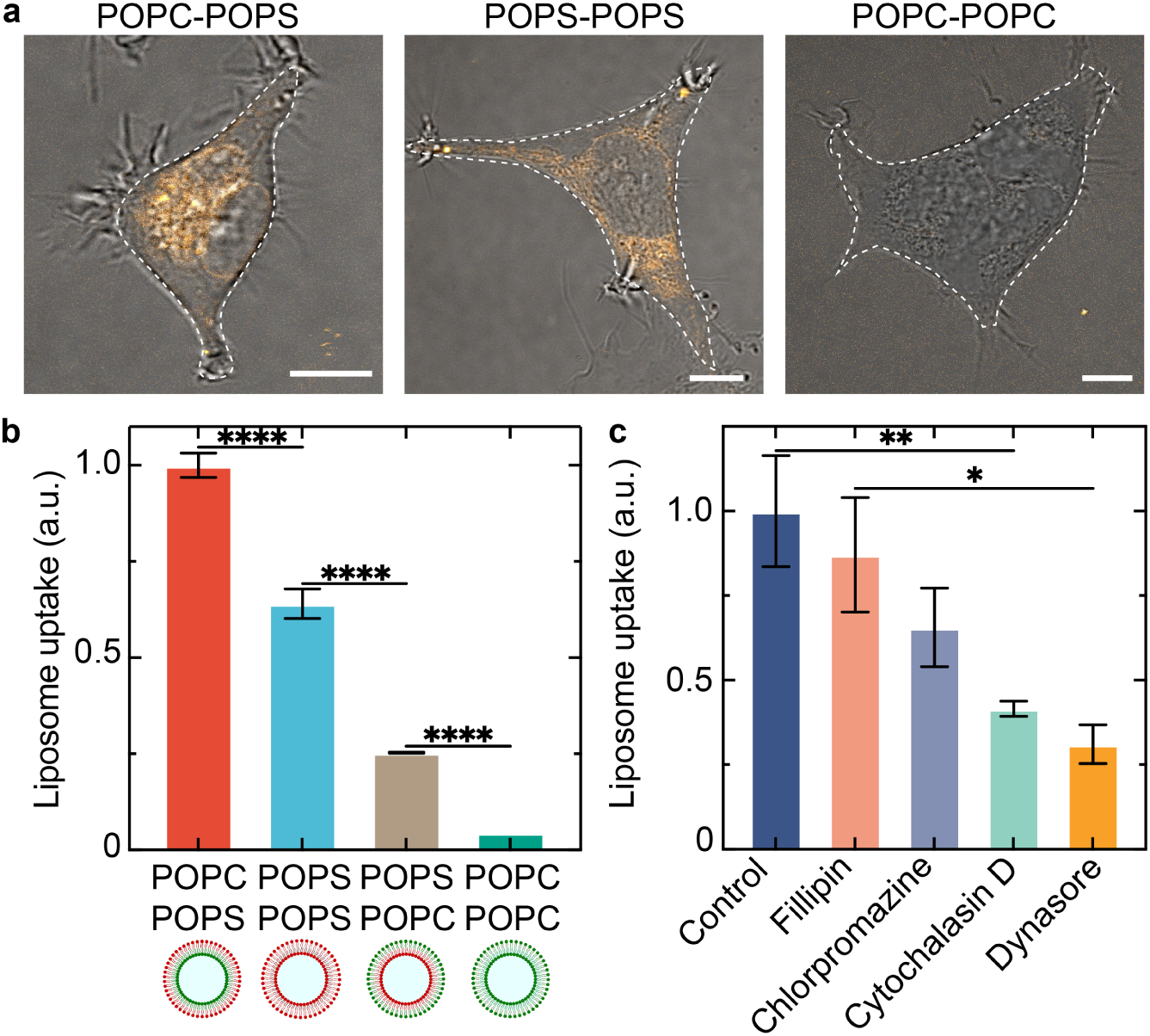
Cell uptake of asymmetric lipid vesicles. **a** Confocal images of HEK293 cells after incubation with POPC-POPC, POPS-POPS and POPC-POPS vesicles for 6 h (orange, labeled with Rhodamine B, (*λ_ex_*=561 nm). Scale bars: 10 µm. **b** Lipid vesicle uptake of cells analyzed with flow cytometry (n*>*15300, Mean *±* SEM). **c** Lipid vesicle uptake of cells with flow cytometry in the presence of small molecule endocytosis inhibitors. Mean *±* SEM, n>20800.

To investigate if the lipid vesicles can also release cargo into the cytoplasm of cells through endosomal escape, we load them with an mRNA that encodes the green fluorescent protein (GFP). Upon uptake of lipid vesicles by the cells, the cargo ideally releases from the lipid vesicle and escapes from the endosome leading to the translation of the mRNA into GFP as schematized in Figure 4**a**. Upon encapsulating GFP mRNA into POPC-POPS, POPS-POPS, and POPC-POPC lipid vesicles and adding them to HEK cells, we observe the mRNA transfection after 48 hours via confocal imaging. We observe that cells are successfully transfected with GFP as shown in Figure 4**b**, indicating that a fraction of the cargo can escape the endosome. We determine the mRNA transfection efficiency by calculating the percentage of fluorescent cells relative to the total cell number. The transfection efficiency of POPC-POPS vesicles is 9 times higher than POPC-POPC and 7 times higher than POPS-POPS, as shown in Figure 4**c**. We also observe that the intensity of the GFP signal is significantly higher for POPC-POPS compared to the other conditions, indicating that more mRNA has been released into the cytoplasm as shown in Figure S11. These results show that asymmetric vesicles are taken up more than symmetric vesicles and that they are more efficient in undergoing endosomal escape.

**Figure 4:**
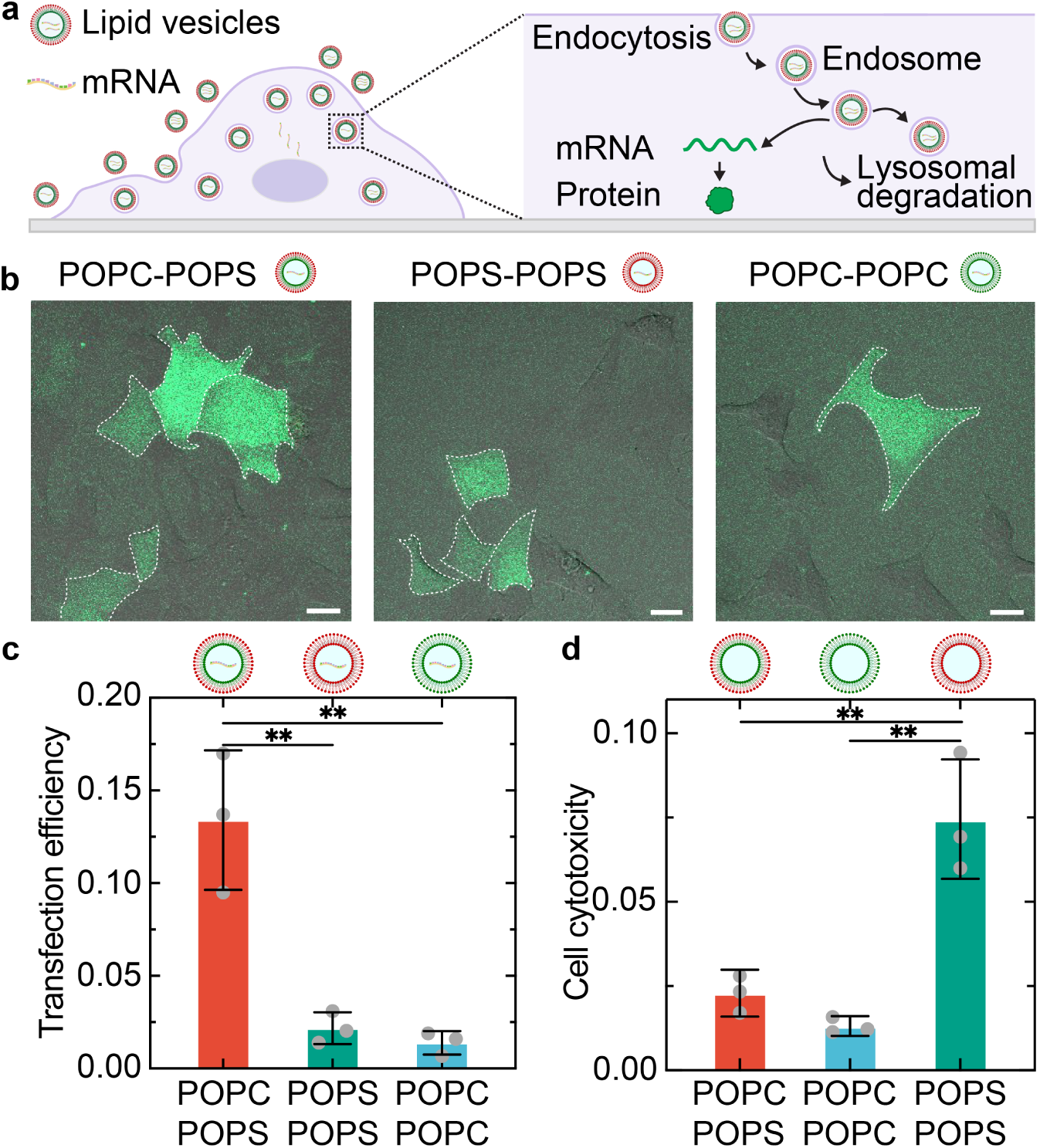
Asymmetric lipid vesicles increase transfection efficiency of HEK293 cells. **a** Schematic representation of the lipid vesicle uptake and mRNA delivery to HEK293 cells. **b** Confocal overlay images of HEK293 cells transfected with GFP (green, *λ_ex_*=488 nm) after incubation with POPC-POPC, POPS-POPS and POPC-POPS vesicles. Scale bars: 20 µm. **c** mRNA transfection efficiency measured as ratio of transfected cells. **d** Cell cytotoxicity of asymmetric vesicles measured with an LDH-assay. Mean *±* SD, n=3.

Cell membranes are asymmetric, with a precise positioning of lipids in the inner or outer leaflet. Conventional symmetric lipid vesicles that are taken up by cells can cause a disruption of the cell membrane and lipidome leading to cell apoptosis.^32^ To investigate, if asymmetric lipid vesicles that resemble the structure of cell membranes, could reduce the cell cytotoxicity, we employ a lactate dehydrogenase (LDH) assay to measure the cell viability after lipid vesicle addition. We find that the cytotoxicity of POPC-POPS is significantly lower than that of POPS-POPS despite its 2-fold higher uptake by cells, as depicted in Figure 4**d**. Thus, asymmetric lipid vesicles can result in a reduced cell cytotoxicity likely because their membrane resembles the cell membrane, which contains PS-lipids on the inner leaflet. Overall, the excellent properties of asymmetric lipid vesicles revealing an increased cell uptake efficiency, enhanced mRNA transfection and lower toxicity make them an efficient and unique drug delivery carrier that offers full programmability over their biochemical and biophysical properties.

The delivery of proteins is an important strategy for disease treatment because proteins can directly manipulate cellular functions. Compared to the delivery of mRNA, proteins provide direct functionality once they are released in cell cytoplasm, resulting in a more efficient and rapid treatment for diseases. To further explore the versatility of the asymmetric lipid vesicles as delivery vehicles beyond mRNA, we encapsulate different fluorescently labeled proteins and add the asymmetric vesicles to HEK cells. The confocal images in Figure 5**a** reveal the presence of proteins inside cells. These proteins have very different functions and molecular weights ranging from 60 kDa for streptavidin to 240 kDa for B-phycoeryhtrin. By comparing the signal of lipid vesicles and proteins, we also find that a fraction of proteins has been released into the cytoplasm because their signal is not colocalized with the lipid vesicles. We also verify protein functionality by using a Cas9 labeled with GFP; the observed GFP fluorescence confirms that the protein is functional. Strikingly, we even observe that GFP-Cas9 gradually targets the nucleus within 6 hours, due to the presence of nuclear localization signals on the protein surface. By analyzing the fluorescence intensity of GFP-Cas9 inside the cell, we find that almost 40% of the Cas9 is localized in the nucleus, as shown in Figure 5**b**. This further shows that the Cas9 protein is released from asymmetric lipid vesicles and remains functional.

**Figure 5:**
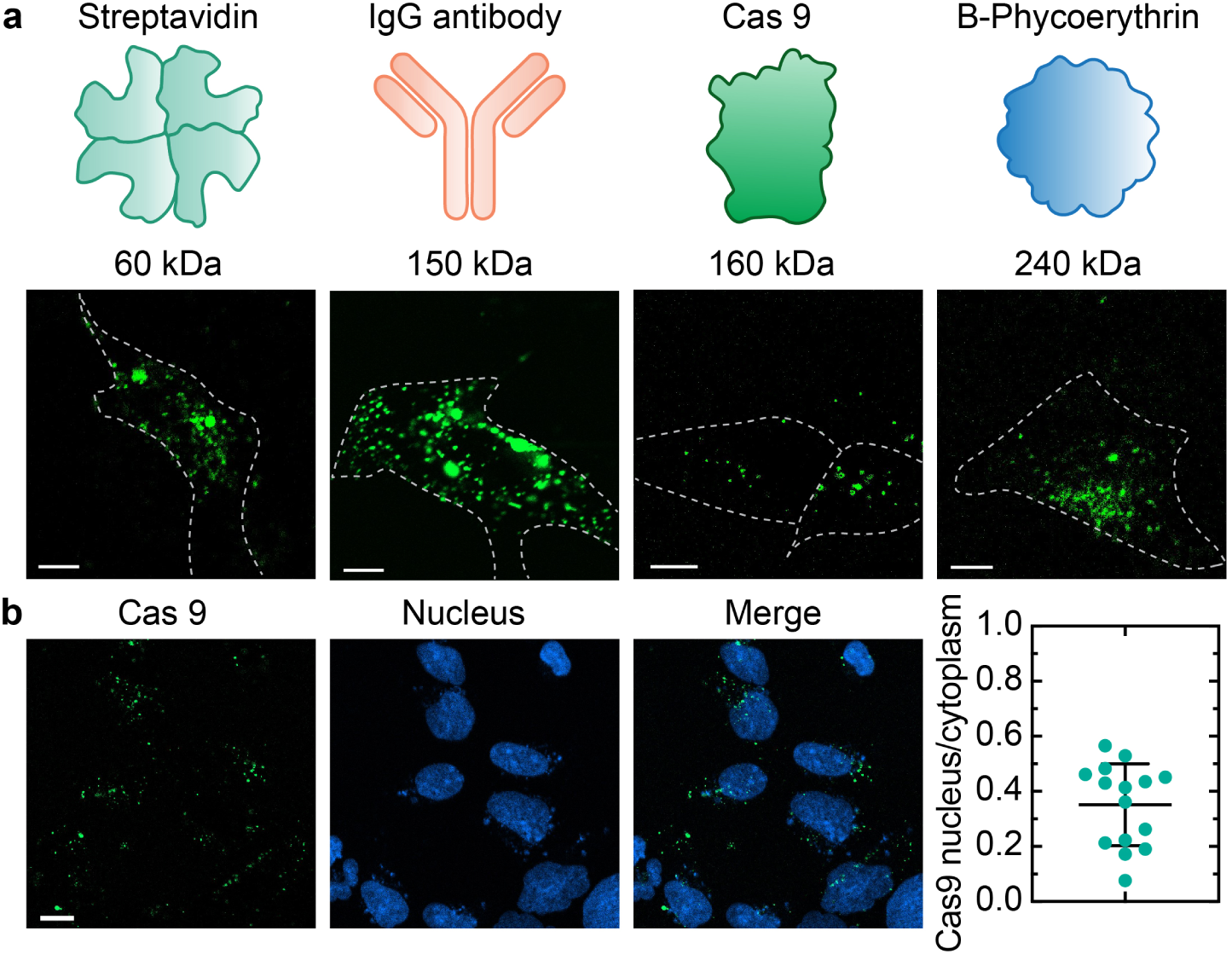
Asymmetric lipid vesicles deliver proteins to cells. **a** Cytosolic delivery of fluorescent proteins to HEK293 cells. Scale bars: 20 µm. (Streptavidin: *λ_ex_*=647 nm; IgG: *λ_ex_*=488 nm; Cas 9: *λ_ex_*=488 nm, B-Phycoerythrin: *λ_ex_*=561 nm) **b** The cytoplasmic release and nucleus targeting of a GFP-fused Cas9 protein. Scale bar: 10 µm. (GFP: green, *λ_ex_*=488 nm, Hoechst: blue, *λ_ex_*=455 nm). Mean *±* SD, n=15.

## Perspective

The engineering of asymmetric lipid vesicles with distinct lipid compositions in the inner and outer leaflet enables the design of new types of delivery vehicles. Our method allows the screening of large libraries of lipids and the generation of lipid vesicles with properties that could not be assessed before. It also prompts to investigate membrane biophysics and biochemistry with full control over the positioning of individual lipids. It will be particularly interesting to identify the exact molecular mechanisms that lead to a reduction of the cell cytotoxicity when using asymmetric vesicles. In comparison to symmetric lipid vesicles, our findings show that an asymmetric leaflet configuration enhances the cellular lipid vesicle uptake, improves the mRNA transfection efficiency, and enables the effective delivery of functional proteins. Thus, the ability to design asymmetric lipid vesicles on the nanoscale opens important applications for drug delivery ranging from vaccines to cancer treatment. We have shown that we can separately deliver Cas9 and RNA using asymmetric vesicles. Thus, our method could potentially also be used to deliver a ribonucleoprotein complex containing Cas9 and a single guide RNA to enable the full gene editing potential of Cas9.

## Experimental Section

### Emulsion preparation and extrusion

To form asymmetric lipid vesicles, we prepared two solutions of mineral oil (Sigma Aldrich): one for the inner leaflet and one for the outer leaflet.^33^ Each solution contained a concentration of 10 mg/mL of the respective lipid. The lipids used in this work are 1-palmitoyl-2-oleoyl-sn-glycero-3-phospho-L-serine (POPS), 1-palmitoyl-2-oleoyl-glycero-3-phosphocholine (POPC), 1,2-dioleyloxy-3-dimethylaminopropane (DODMA), 1-palmitoyl-2-oleoyl-sn-glycero-3-ethylphosphocholine (EPC), 1,2-dioleoyl-sn-glycero-3-phosphoethanolamine-N-(lissamine rhodamine B sulfonyl) (LissRhod-PE), 1,2-dioleoyl-sn-glycero-3-phosphoethanolamine-N-(7-nitro-2-1,3-benzoxadiazol-4-yl) (NBD-PE). Lipids were purchased from Avanti Polar Inc., dissolved in chloroform and stored at *−*20 *^◦^*C until use. Lipids were dried under a stream of nitrogen and resuspended in mineral oil. The lipid composition contained 99 mol% of POPS, POPC or DODMA, respectively, and 1 mol% LissRhod-PE, if not stated otherwise. To facilitate the resuspension, the mineral oil solutions were sonicated and incubated for 60 to 90 min. Then 700 µL of the second mineral oillipid solution was layered on top of 600 µL 1x PBS and also left to incubate for 2 h. Then lipid-stabilized emulsions were formed by mixing 10 µL 1x PBS (Gibco) with 500 µL mineral oil/lipid solution. The mixture was briefly vortexed to from water-in-oil droplets. Subsequently, the droplets were extruded nine times (Avanti Polar) through a polycarbonate membrane with a pore size of 200 nm (unless otherwise specified). Then, 100 µL of the droplets were added on top of the second oil solution and centrifuged for 10 min at 10 000 g. Due to the density gradient between water and mineral oil, the lipid-stabilized emulsions were centrifuged through the lipid monolayer at the oil-water interface and dissolved in the aqueous phase. Finally, the aqueous phase containing asymmetric vesicles was pipetted out with a syringe needle (Air-Tite) and stored in LoBind tubes (Eppendorf) for up to one day. Typically, vesicles were used immediately after their formation.

### Dynamic light scattering

The hydrodynamic diameter of lipid vesicles was determined with a ZetaSizer Nano ZS instrument (Malvern Pananalytical) that utilizes a He–Ne laser with a wavelength of 632 nm and applies the phase analysis light scattering technique. Lipid vesicles were diluted 1:10 for corresponding hydrodynamic diameter measurements by placing 100 µL of dispersion in ZEN0040 disposable plastic microcuvettes. The size measurement results from DLS measurements are displayed as an intensity weighted distribution function of particle size, with determined intensity weighted Z average (mean size) and the polydispersity index (PDI).

The zeta potential of lipid vesicles was determined using a Zetasizer Pro (Malvern Panalytical), which uses electrophoretic light scattering to measure the velocity of particles under an applied electric field, providing information about their surface charge. For each measurement, 1 mL of lipid vesicle dispersion was placed into a DTS1070 disposable folded capillary cell, and an electric field of 50 to 100 mV was applied to induce particle movement. Each sample was measured three times, with each measurement being the average of 20 individual readings. The zeta potential values were calculated from the electrophoretic mobility and are reported in millivolts (mV).

### Quenching assay

The leaflet asymmetry was measured using fluorescence quenching experiments. For these, 1 mol% fluorescent lipid tracer molecule (NBD-PE, Avanti) was added to the outer leaflet. After vesicle formation, 100 µL of vesicles were placed in a well and its fluorescence measured every minute with a microplate reader. The excitation wavelength was set to 465 nm and the emission was detected at 536 nm. After three minutes, 10 µL of 200 mM sodium dithionite (Sigma Aldrich) were added to the vesicles. Dithionite quenches NBD-fluorophores and leads to a reduction of fluorescence intensity, when NBD is present in the outer leaflet. NBD on the inside of lipid vesicles is not quenched. After 15 min, the quenching reaction saturated. To prove that the remaining fluorescence originates from NBD-PE in the inner leaflet of lipid vesicles, 5 µL of Triton X were added to disrupt the lipid vesicle stability and expose the remaining NBD to dithionite quenching.

### Pendant drop

The interfacial tension between mineral oil and 1x PBS was measured using the pendant drop method (Droplet Lab). The respective lipid was dissolved in mineral oil at a concentration of 0.06 mM. Since mineral oil has a lower density than water, an oil droplet was pushed into the aqueous solution using a syringe needle. The trace of the oil droplet was detected and analyzed with a Young-Laplace fit to determine the interfacial tension.

### Partitioning assay

Rhodamine 6G (Sigma Aldrich) was dissolved to a concentration of 1 mM in 1x PBS. To study the partitioning of lipids and micelle formation, 1 ml of this solution was placed in a cuvette and 1 ml of the respective mineral oil/lipid solution was layered on top. The cuvettes were imaged after 0, 12, 24, and 36 h. In addition, 50 µL of the aqueous solution was removed at each time point and placed in a 96-well plate. Its absorbance and fluorescence emsission spectrum after excitation with 500 nm measured.

### Determination of vesicle concentration

The vesicle concentration was measured using 1 mol% of a rhodamine-labeled fluorescent lipid in the lipid composition. Following the centrifugation step, 100 µL of the vesicle solution was placed in a 96-well plate. The fluorescence intensity of each well was measured with a microplate reader (Tecan Spark) using an excitation wavelength of 540 nm and detecting the emission at 570 nm. The fluorescence intensity is proportional to the vesicle concentration and was used to normalize the amount of vesicles that was added to cells.

### Cell culture

Human embryonic kidney (HEK) cells were dispersed in 10 mL of Dulbecco’s Modified Eagle Medium (DMEM) media supplemented with 10 % fetal bovine serum and 1 % pennicilin/streptomycin in a 75 mL flask. The cells were kept in an incubator at 37 *^◦^*C and 5% CO_2_. Every 48 hours the media was exchanged by aspirating the media using a vacuum and washing the cells with 10 mL of 1x phosphate buffered saline (PBS) upon which 10 mL of fresh media was added. When the cell density was around 80%, the cells were passaged. The media was removed and 2.5 mL of trypsin is added in the flask for 3 min to detach cells from the bottom of the flask. Then the flask was taken out of the incubator and 5 mL of media are added. The cells suspension was centrifuged for 5 min at 500 g. Once the cells were concentrated at the bottom of the tube, the media and trypsin were aspirated and the cells resuspended in fresh media. The cell concentration was measured before centrifugation to calculate the volume of media that needs to be added at this step to tune the cell concentration used in the uptake and transfection experiments.

### Confocal microscopy

Confocal microscopy was performed on a confocal laser scanning microscope (LSM980 or LSM900) by Zeiss (Carl Zeiss AG). The pinhole was set to 1 airy unit and imaging was performed at 37 *^◦^*C and 5 % CO_2_. For image acquisition a 20x air (Plan-Apochromat 20x/0.8 M27), a 40x water immersion objective (Plan-Apochromat 40x/1.0 DIC M27) and a 63x oil immersion (HC PL APO 63x/1,40 OIL CS2) were used. Images were analyzed and adjusted in ImageJ.

### Lipid vesicle uptake and mRNA transfection

For the cell experiments, 12 or 24 well plates were used with cell concentrations of approximately 1.0*×* 10^5^ cells/mL. The first step of the cell experiments was to seed the cells on day 0 in the wells, then add freshly made lipid vesicles on day 1 and observe them with confocal microscopy on days 2, 3 and 4 after having exchanged the media to remove all remaining lipid vesicles that had not been yet taken up by the cells. Before the addition of lipid vesicles, their concentration was measured with the microplate reader. This was used to normalize the amount of vesicles added to each well. Typically, we added 100-200 µL of lipid vesicles to each well. To measure the amount of lipid vesicle uptake per cell, the contour of 20 cells per image was determined using ImageJ. Then the mean vesicle fluorescence intensity inside each cell is measured. The mean for the 20 cells per well as the SD is then calculated for each condition.

### mRNA delivery

To measure the cell transfection efficiency, 2 µL GFP-encoding mRNA was added to the oil phase before the extrusion of emulsion droplets. Confocal images were taken on day 1, 2 and 3. For each well, multiple images are taken and for each image the following ratio was determined measured: *N*_transfected_*/*(*N*_transfected_ + *N*_untransfected_). Multiple ratios per well were obtained to calculate the mean transfection ratio and standard deviation for each condition. Complementing the GFP fluorescence, LissRhod PE was used as fluorescent lipid.

### Protein delivery

We delivered various proteins, such as Immunoglobulin GAlexa Fluor 488, Jackson ImmunoResearch, Streptavidin (Streptavidin, Alexa Fluor 647 conjugate, ThermoFisher Scientific), B-Phycoerythrin (Sigma Aldrich), and GFP Cas9 (Integrated DNA Technologies) using asymmetric lipid vesicles to HEK 293 cells. To achieve this, 2 µL of protein solution of different proteins were added to the oil phase before the extrusion of emulsion droplets. Confocal images were taken on days 1, 2 and 3. Typically, after 48 h most proteins were degraded.

### Flow cytometry

Following the lipid vesicle uptake, cells were trypsinized and concentrated to 10 *×* 10^6^ cell*/*mL. Flow cytometry was performed using a BD FACSymphony A3 Lite (BD Biosciences) instrument with laser wavelength 561 nm. The data was analyzed with FCS express software. Cells were gated to remove cell debris (FSC-A/SSC-A). For each condition 20000 cells were analyzed.

### Cell cytotoxicity assay

Cell cytotoxicity was determined using a CyQUANT LDH cytotoxicity assay (Thermo Fisher). Following the manufacturers protocol, cells we first measured the LDH activity as a function of the cell number revealing an ideal cell number of 2500 cells*/*well. Cells were seeded in triplicates per condition and the respective lipid vesicles added after 24 h. After another 24 h, the spontaneous, maximum and lipid vesicle-induced LDH activity was measured with a plate reader for each condition. The cytotoxicty was then calculated according to: 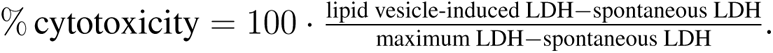

### Statistical Analysis

All the experimental data were reported as mean *±* SD from n experiments. The respective value for n is stated in the corresponding figure captions. All experiments were repeated at least twice. To analyze the significance of the data, a Student’s t-test with Welch’s correction was performed using Prism GraphPad (Version 9.1.2) and p-values correspond to ****: p *≤* 0.0001, ***: p *≤* 0.001, **: p *≤* 0.01, *: p *≤* 0.05 and ns: p *≥* 0.05.

## Acknowledgement

K.J. thanks the Joachim Herz Foundation and the Alexander von Humboldt Foundation for financial support. This work was supported in part by the Harvard MRSEC program of the NSF under Award No. DMR 20-11754 and the HealthInnoHK program of the Innovation and Technology Commission of the Hong Kong SAR Government. The authors thank the Harvard Center for Biological Imaging (RRID:SCR_018673) and the Harvard GRID Accelerator for infrastructure and support. This work was performed in part at the Harvard University Center for Nanoscale Systems (CNS); a member of the National Nanotechnology Coordinated Infrastructure Network (NNCI), which is supported by the National Science Foundation under NSF award no. ECCS-2025158. The authors thank the Bauer Core Facility at Harvard University. We thank Laura Rodriguez Arriaga and Tuomas Knowles for helpful discussions.

## Supporting Information Available

Supporting Information is available.

