## Supporting Information for "Engineering asymmetric nanoscale lipid vesicles for drug delivery"

Supporting Figure 11: GFP fluorescence intensity measured in transfected cells . 14

### Supporting Notes

#### Supporting Note 1: Estimation of the vesicle concentration

To determine the vesicle concentration after the centrifugation step, we calculate the average amount of lipids per vesicle. The surface area of a vesicle is given as:

$$A = 4\pi r^2\,, \tag{1}$$

where $r$ is the vesicle radius, which we determined to 200 nm. To obtain the number of lipids per vesicle $N_l$, we use the lipid headgroup area of 0.7 nm$^2$ and multiply the area to account for the lipid bilayer:

$$N_l = 2A/0.7\text{nm}^2 = 360000\,. \tag{2}$$

To calculate the density of vesicles $N_v$, we use the measured lipid concentration $c$ of 65 µM for POPC-POPS vesicles and insert it into:

$$N_v = c \cdot N_A/N_l = 1.1 \cdot 10^{14}\,\text{vesicles/l} \tag{3}$$

with $N_A$ being the Avogadro constant. Given that our usual sample volume was 500 µL, this leads to the formation of $5.5 \cdot 10^{10}$ vesicles during the experiment. Finally, we calculate the liposome formation efficiency, which is the ratio of the encapsulated volume and the emulsion volume. This is given as:

$$E = \frac{4}{3}\pi r^3 \cdot N_v/V_i = 0.023\,, \tag{4}$$

with $V_i$ being the initial emulsion volume, which was 10 µL. Thus, we achieve an encapsulation efficiency of 2.3 %. Notably, the initial volume can be reduced to achieve an even higher encapsulation efficiency.

### Supporting Figures

#### Supporting Figure 1: TEM images of asymmetric lipid vesicles

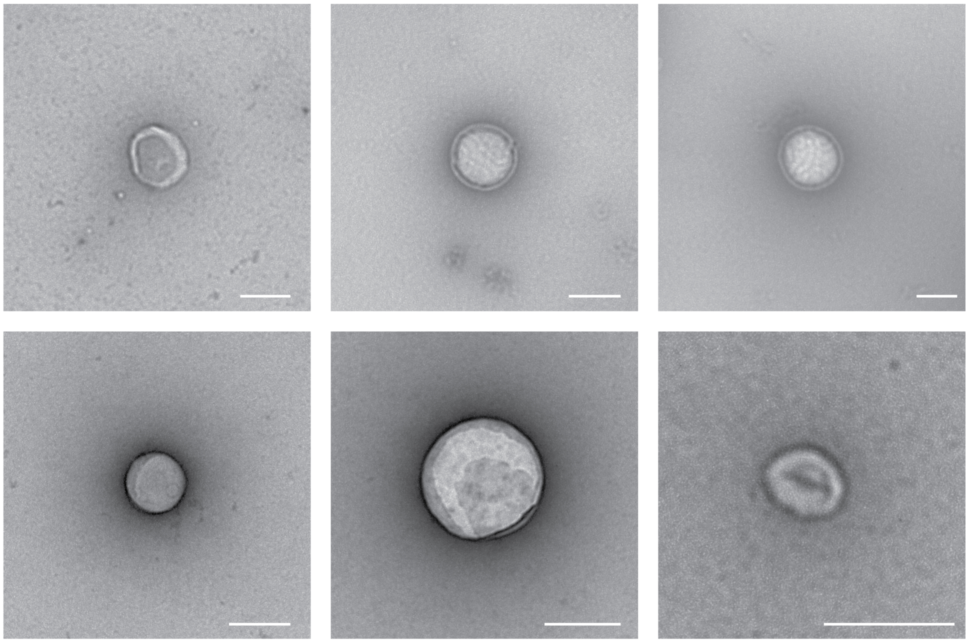

Supporting Figure 1: TEM images of asymmetric lipid vesicles. Scale bars: 200 nm

#### Supporting Figure 2: Dithionite quenching experiment for different lipid compositions

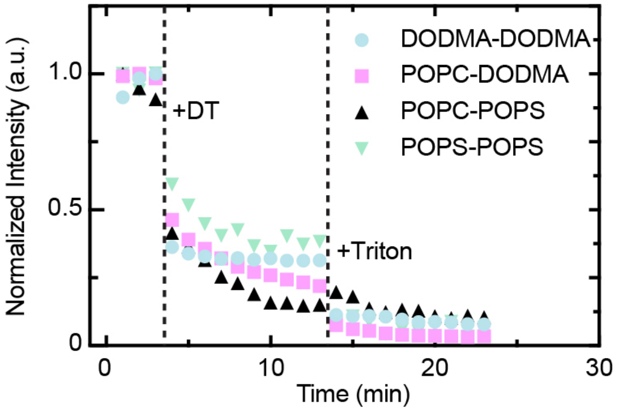

Supporting Figure 2: Dithionite quenching experiment for different lipid compositions.

#### Supporting Figure 3: Zeta potential for different lipid compositions for inverted and extruded lipid vesicles

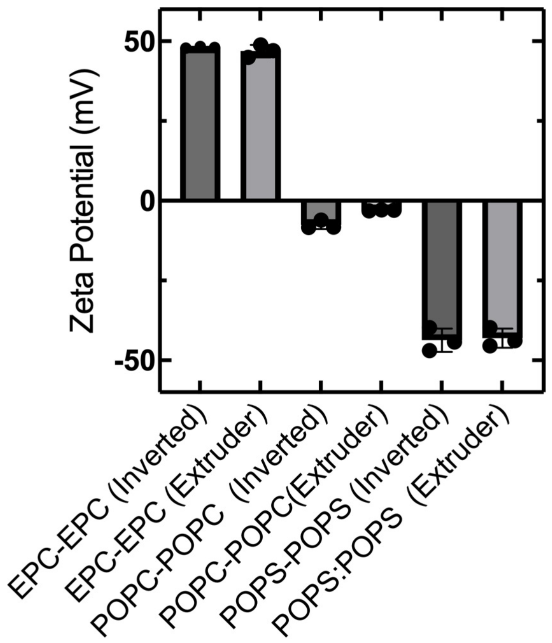

Supporting Figure 3: Zeta potential for different lipid compositions for inverted and extruded lipid vesicles.

#### Supporting Figure 4: Molecular structures of POPC, POPS, DODMA and EPC

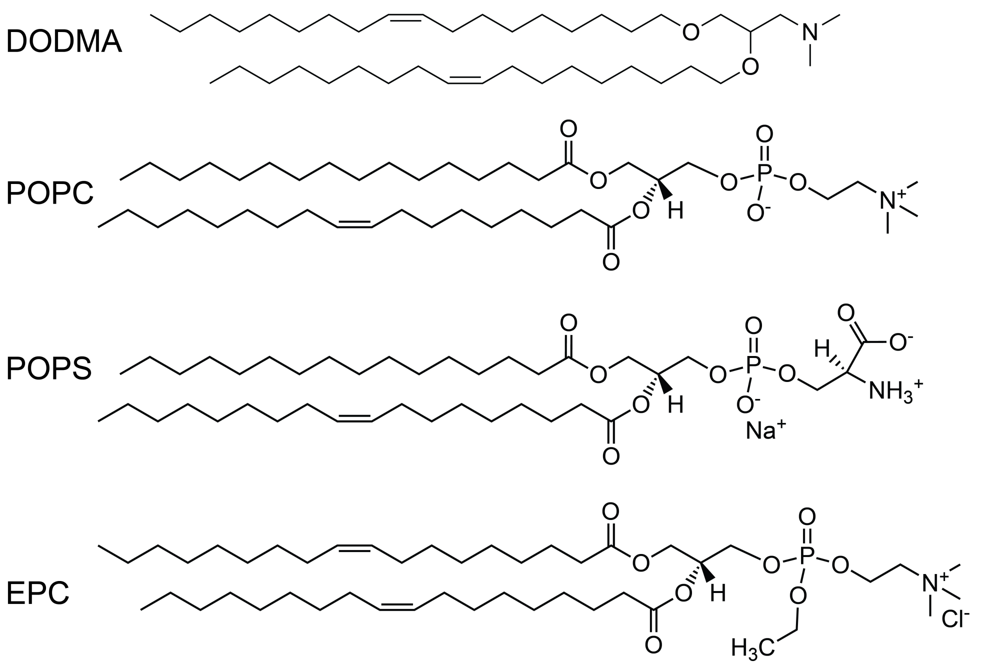

Supporting Figure 4: Molecular structures of POPC, POPS, DODMA and EPC.

#### Supporting Figure 5: Partitioning experiments with Rhodamine 6G

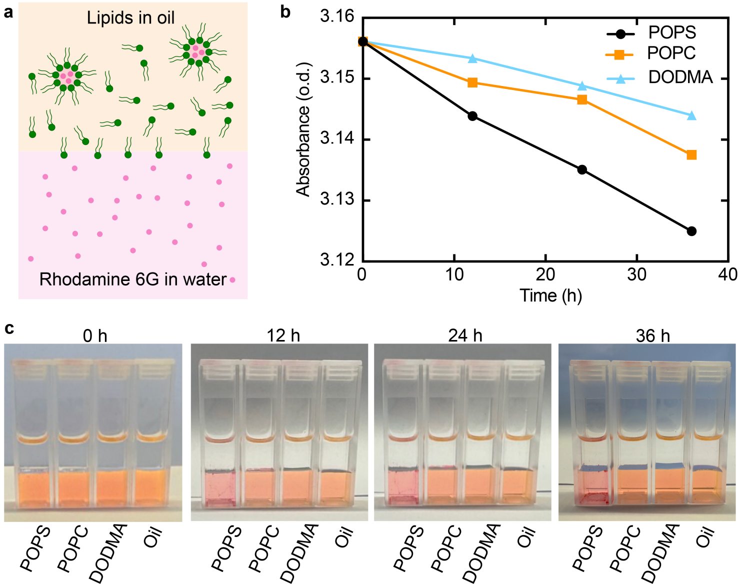

Supporting Figure 5: Partitioning experiments of Rhodamine 6G. Aqueous solutions of Rhodamine 6G 1 mM (bottom phase) are exposed to the oil phase containing different lipids concentrations in a 3.5 ml cuvette. **a** Schematic representation of the extraction of Rhodamine 6G from the aqueous to the oil phase due to inverted micelle formation of lipid molecules that enclose rhodamine 6G molecules. **b** Rhodamine 6G absorbance ($\lambda_{ex} = 500\,\text{nm}$)in the aqueous solution over time partitioning experiment.**c**Optical images of the variation of Rhodamine aqueous solution over time.

#### Supporting Figure 6: Calibration of rhodamine-labeled lipids

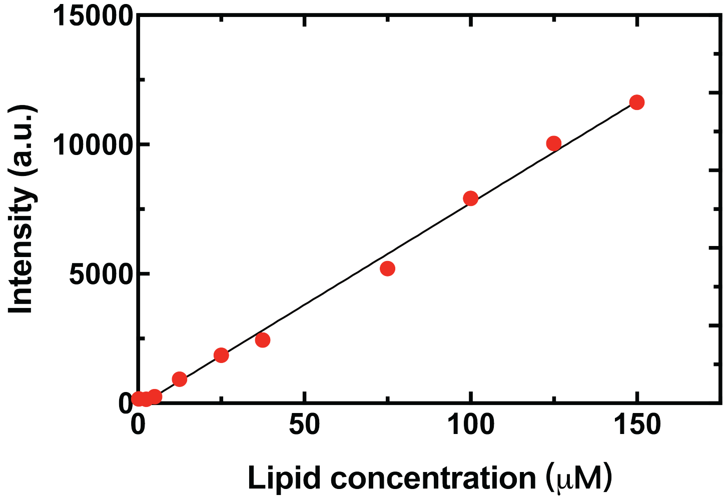

Supporting Figure 6: The relationship between fluorescent intensity of Lissamine Rhodamine and lipid concentration.

#### Supporting Figure 7: Cell uptake of different symmetric liposomes formed by extrusion

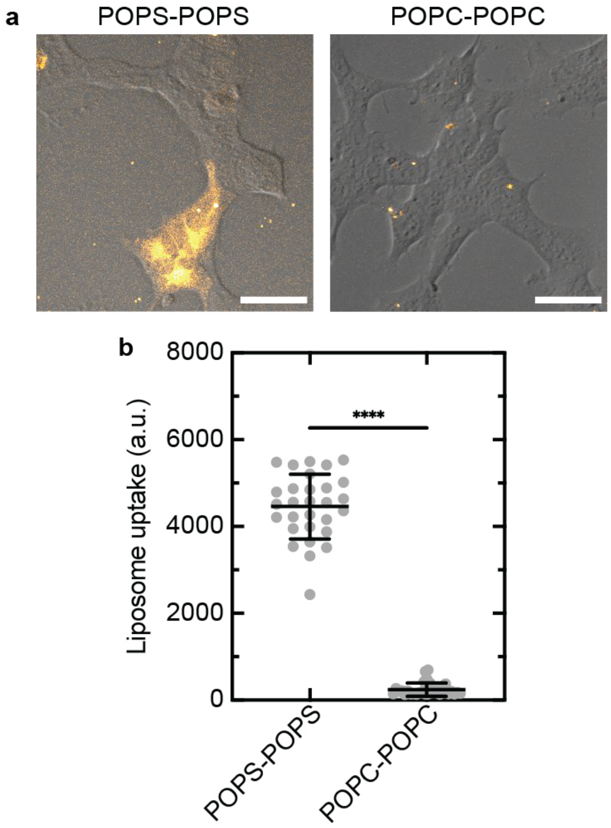

Supporting Figure 7: Cell uptake of different symmetric liposomes by extrusion. **a** Confocal images of HEK293 cells after incubation with liposomes for 24 h (orange, labeled with Rhodamine B, ($\lambda_{ex}$ = 561 nm, n>20, Mean ± SD) Scale bars: 10 µm. **b** Liposome uptake of cells analyzed from confocal images.

#### Supporting Figure 8: Cell uptake of asymmetric liposomes

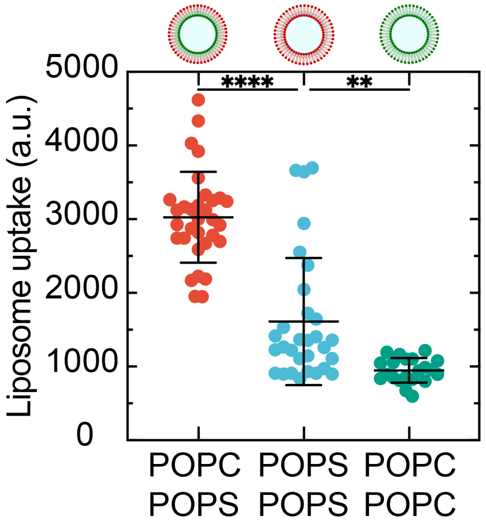

Supporting Figure 8: Cell uptake of asymmetric liposomes quantified with confocal microscopy.

#### Supporting Figure 9: Liposome uptake of POPC-POPS with the presence of small molecules endocytosis inhibitors

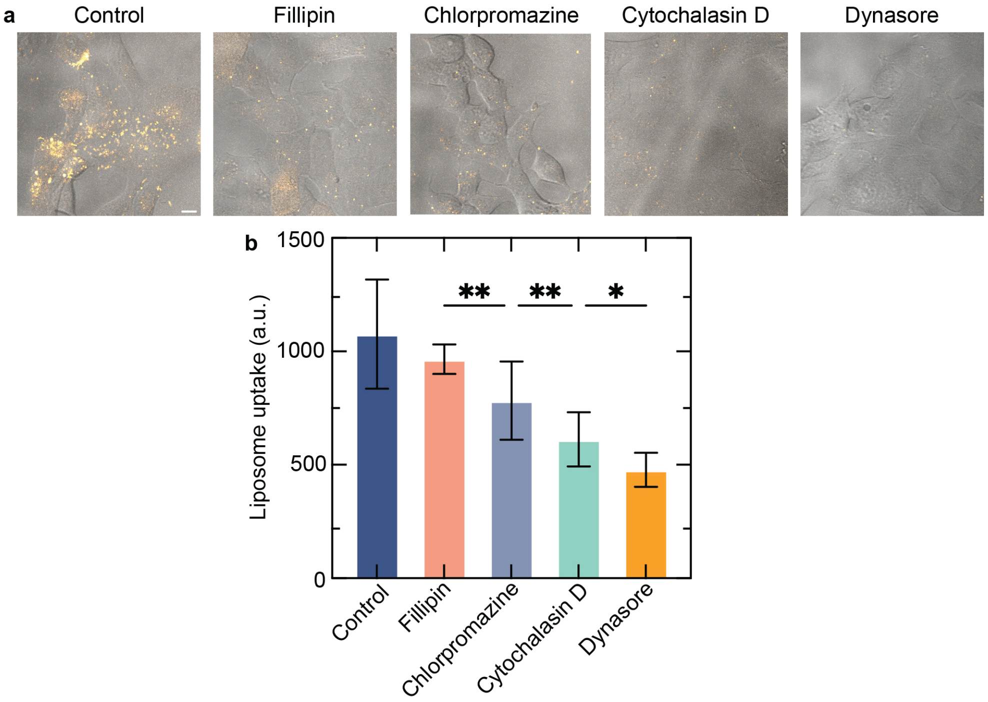

Supporting Figure 9: Liposome uptake of POPC-POPS with the presence of small molecules endocytosis inhibitors. **a** Confocal images of HEK293 cells after incubation with liposomes for 4 h (orange, labeled with Rhodamine B, ($\lambda_{ex} = 561\,\text{nm}$). Scale bars: 10 µm. **b** Liposome uptake of cells analyzed from confocal images.

#### Supporting Figure 10: Liposome uptake for different fatty acid lengths

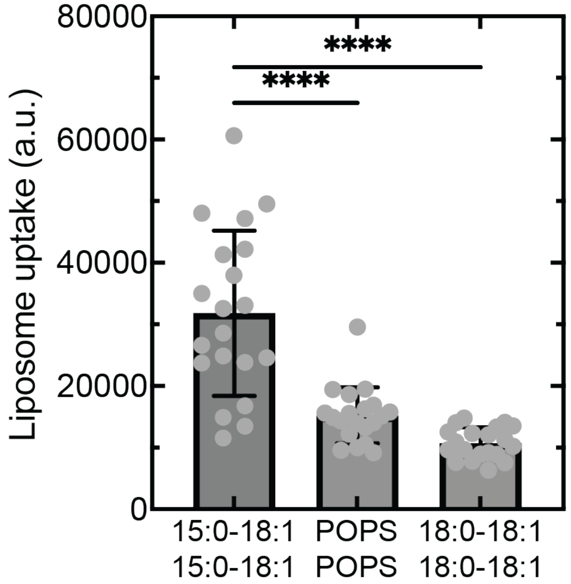

Supporting Figure 10: Liposome uptake of symmetric vesicles with different length of the fatty acids from 15:0 PS to 18:0 PS.

#### Supporting Figure 11: GFP fluorescence intensity measured in transfected cells

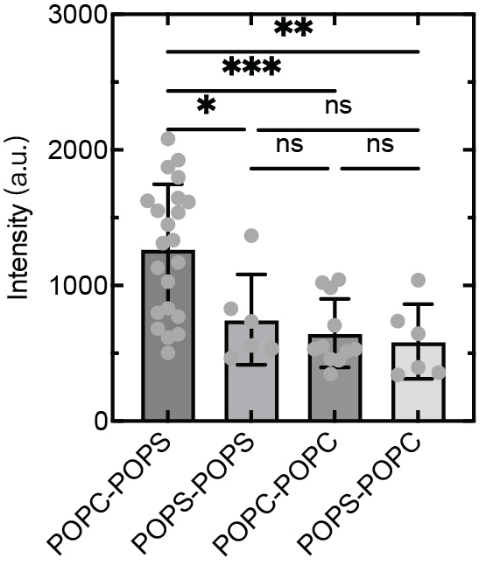

Supporting Figure 11: GFP fluorescence intensity measured in transfected cells.
